# Germline loss-of-function variants in the base-excision repair gene *MBD4* cause a Mendelian recessive syndrome of adenomatous colorectal polyposis and acute myeloid leukaemia

**DOI:** 10.1101/2021.04.27.441137

**Authors:** Claire Palles, Edward Chew, Judith E. Grolleman, Sara Galavotti, Christoffer Flensburg, Erik A.M. Jansen, Helen Curley, Laura Chegwidden, Edward Arbe Barnes, Ashish Bajel, Kitty Sherwood, Lynn Martin, Huw Thomas, Demetra Georgiou, Florentia Fostira, Yael Goldberg, David J. Adams, Simone A.M. van der Biezen, Michael Christie, Mark Clendenning, Constantinos Deltas, Aleksandar J. Dimovski, Dagmara Dymerska, Jan Lubinski, Khalid Mahmood, Rachel S. van der Post, Mathijs Sanders, Jürgen Weitz, Jenny C. Taylor, Clare Turnbull, Lilian Vreede, Tom van Wezel, Celina Whalley, Claudia Arnedo Pac, Gulio Caravagna, William Cross, Daniel Chubb, Anna Frangou, Andreas Gruber, Ben Kinnersley, Boris Noyvert, David Church, Trevor Graham, Richard Houlston, Nuria Lopez-Bigas, Andrea Sottoriva, David Wedge, Genomics England Research Consortium, The CORGI Consortium, WGS500 Consortium, Mark A. Jenkins, Roland P. Kuiper, Andrew W. Roberts, Marjolijn J.L. Ligtenberg, Nicoline Hoogerbrugge, Viktor H. Koelzer, Andres Dacal Rivas, Ingrid M. Winship, Clara Ruiz Ponte, Daniel D. Buchanan, Ian P.M. Tomlinson, Ian J. Majewski, Richarda M. de Voer

**Author notes:** **Authors for correspondence**: Ian P. M. Tomlinson -, Ian J. Majewski -, Richarda M. de Voer –. See appendix (Genomics England Research Consortium, The CORGI Consortium, WGS500 Consortium). These authors contributed equally (Claire Palles, Edward Chew, Judith E. Grolleman, Sara Galavotti, Christoffer Flensburg). These authors jointly supervised this project (Ian P.M. Tomlinson, Ian J. Majewski, Richarda M. de Voer).

## Abstract

Inherited defects in base-excision repair (BER) predispose to adenomatous polyposis and colorectal cancer (CRC), yet our understanding of this important DNA repair pathway remains incomplete. By combining detailed clinical, histological and molecular profiling, we reveal biallelic germline loss-of-function (LOF) variants in the BER gene *MBD4* to predispose to adenomatous polyposis and –uniquely amongst CRC predisposition syndromes– to myeloid neoplasms. Neoplasms from MBD4-deficient patients almost exclusively accumulate somatic CpG>TpG mutations, resembling mutational signature SBS1. MBD4-deficient adenomas harbour mutations in known CRC driver genes, although *AMER1* mutations were more common and *KRAS* mutations less frequent. We did not find an increased risk for colorectal tumours in individuals with a monoallelic *MBD4* LOF variant. We suggest that this condition should be termed *MBD4*-associated neoplasia syndrome (MANS) and that *MBD4* is included in testing for the genetic diagnosis of polyposis and/or early-onset AML.

## INTRODUCTION

Several syndromes characterized by adenomatous polyposis and colorectal cancer (CRC) are caused by high-penetrance germline pathogenic variants that directly impair the correction of mispaired bases within DNA. These syndromes include the dominantly inherited condition polymerase proofreading-associated polyposis (PPAP) caused by pathogenic variants in the polymerase proofreading domains of *POLE* and *POLD1*,^1^ and the recessively inherited conditions constitutional mismatch-repair deficiency, caused by variants in genes involved in mismatch repair (*MSH2, MLH1, PMS2, MSH6*),^2,3^ and *MUTYH*-associated polyposis and *NTHL1*-associated tumour syndrome, caused by genes involved in base excision repair (BER) (*MUTYH* and *NTHL1*).^4,5^ In each case, there are also increased risks of specific extra-colonic tumours. Although the mechanistic connection between these germline variants and colorectal tumours is not entirely clear, it is likely that the resulting increased somatic mutation rate in turn causes more somatic mutations in CRC driver genes such as *APC, KRAS* and *TP53*. Previous studies have linked the defect in each DNA repair gene to specific mutational signatures in tumours that align with their repair activity.^6-9^

Despite the identification of roughly 15 colorectal polyposis predisposition genes to date, genetic testing currently fails to identify a cause for a significant proportion of patients who develop multiple colorectal adenomas. Whilst some of these cases may have polygenic or non-genetic origins, it is important to identify any remaining high-risk polyposis predisposition genes in order to prevent CRC in probands and their relatives. Here, we used whole-genome sequencing (WGS) to identify the BER gene *MBD4* as a novel colorectal polyposis predisposition gene with recessive inheritance. By molecular profiling we describe how biallelic inactivating germline variants in *MBD4* predispose to both colorectal polyposis and AML. Furthermore, our data suggest that individuals with monoallelic loss-of-function (LOF) variants in *MBD4* are not at increased risk of polyposis and/or CRC.

## METHODS

### Patient and control cohorts

i. *Discovery cohort:* Oxford-Illumina WGS500 project: constitutional DNA from 35 probands from the UK CORGI study with at least 10 colonic adenomas before age 60 were whole-genome sequenced as previously described.^1^
ii. *Replication cohorts:*
  a. Polyposis and AML: 13 unrelated patients with synchronous or metachronous colorectal polyposis and/or CRC and myelodysplastic syndrome (MDS) and/or AML. Samples were screened for *MBD4* using molecular inversion probes (MIP, *n=*11, **Supplemental Methods**) or whole-exome sequencing (WES; *n=*2). Another 13 cases with synchronous or metachronous colorectal polyposis and/or CRC and MDS and/or AML were identified in the UKBiobank (see below).
  b. Polyposis and/or CRC: 1,493 patients with unexplained polyposis, familial and/or young CRC and >10 adenomatous polyps (with or without CRC), or CRC in combination with other tumour(s) were recruited. This study was approved by the local medical ethics committee (CMO; study number 2015/2172 of the Radboudumc Nijmegen). All samples were screened using MIP-sequencing **(Supplemental Methods; Supplementary Table 1)**.
  c. Eighty-seven families comprised of 198 CRC and early-onset polyp affected individuals were selected for WGS/WES from participants of the Australasian Colorectal Cancer Family Registry (ACCFR): 1) met the definition of Familial Colorectal Cancer Type*x*(FCCTX) (*n=*55); 2) Amsterdam II clinical criteria (AMII) with two of the defining triad being CRC-affected (*n=*4); or 3) >2 CRC-affected blood relatives family members within families in the same blood line but not meeting FCCTX or AMII criteria (MCF)(*n=*28).
iii. *Population cohorts:*
  a. UKBiobank: we searched the WGS/WES data available for 200,000 participants in UKBiobank (October 26th 2020 release) for germline coding or splicing variants in *MBD4*. This research was conducted under UKBiobank application number 8508.
  b. UK 100,000 Genomes Project (100KGP): we searched the WGS data available (ISAAC pipeline) for 17,243 Caucasian participants of the rare diseases programme (v6 release, participants selected for phenotypes with no increased risk of cancer), 2,438 Caucasian CRC patients included in the cancer programme (v8 release) and 283 Caucasian multiple bowel polyps patients (143 v6 rare diseases, 140 pilot project) for germline coding or splicing variants in *MBD4*.
  c. GnomAD: we searched the full and non-Finnish European populations of the gnomAD database (v2.1) for germline coding or splice site variants in *MBD4*.

### Identification of germline variants

WGS of constitutional DNA was performed using the Illumina HiSeq platform as described previously.^1^ Targeted *MBD4* mutation screening was performed (i) using a custom MIP capture panel and sequenced on a NextSeq500 (Illumina) system, or (ii) using Sanger sequencing based on custom PCR amplicons. Variants predicted to result in a premature stop codon (nonsense or frameshifts) or alternative splicing/splice site defect were classed as LOF variants (details provided in the **Supplemental Methods**). All *MBD4* mutations were validated using independent Sanger sequencing reactions.

### Tumour sequencing and somatic variant calling

Tumour samples and their origins from fresh-frozen or formalin fixed paraffin embedded (FFPE) tissue are listed in **Supplementary Table 2**. In brief, we used the i) Agilent SureSelectXT Human All Exon V6 (Agilent Technologies), ii) Agilent SureSelect Clinical Research Exome version 2 (Agilent Technologies) or iii) Illumina TruSeq exome (Illumina) enrichment kits to produce whole-exome libraries that were sequenced using a NextSeq500 (Illumina) or NovaSeq 6000 (Illumina) **(Supplementary Table 2)**.

Somatic variants were called using SuperFreq (v1.3.2),^10^ Strelka (v2.9.2) and/or Mutect2 (GATK v4.1.0.0) with matched germline samples **(Supplementary Table 2)**. Somatic mutations were annotated as described previously.^7^ Full details of the somatic variant calling, removal of FFPE artefacts and validations are provided in the **Supplementary Methods**.

BAM files from previously published multi-region WES data (generated using the SeqCapEx Exome Enrichment Kit v3.0 (Nimblegen/Roche)) from nine sporadic fresh-frozen adenomas were available **(Supplementary Table 2)**.^11^ Somatic variants were re-called using Mutect2 as above.

### Somatic mutation analysis

For each adenoma, high confidence somatic mutations were identified as described previously with minor modifications.^7^ In brief, somatic variants covered by ≥15x sequencing reads, ≥10% variant allele frequency, and ≥6 variant reads, and (for variants called by Mutect2) with ≥2 variant reads per read pair (to exclude FFPE artefacts) were included. WEHI-2 received an allogeneic bone marrow transplant and variants contributed by the haematopoietic stem cell donor were removed based on clonal tracking in SuperFreq.^10^ For somatic variants identified by SuperFreq, a mean read depth >30 across all samples was required and excluded variants if they were detected above 3% VAF (supported by at least 2 reads) in a sample where the clone was deemed absent (clonality <1%). A representative set of somatic mutations were validated by Sanger sequencing or by processing micro-dissected material from the adenomas with the TruSight Tumor 26 Kit (Illumina).

For each adenoma the number of somatic mutations per Mbp was calculated, with the total capture interval estimated using bases with ≥15 read depth. The mutation spectrum and the number of CpG>TpG transitions was determined. A linear model describing the number of CpG>TpG transitions as a function of age was analysed in base R. Methylation status of the sites with somatic mutation was assessed in public whole genome bisulfite sequencing (WGBS) data from normal sigmoid colon from the Roadmap Epigenomics Consortium et al. (as per^12^). The contribution of mutational signatures to the somatic mutation spectrum was inferred using the R package MutationalPatterns^13^ in combination with COSMIC-v3 mutational signatures.^14,15^

All non-synonymous somatic variants in cancer driver genes reported by The Cancer Gene Census (CGC v92) in COSMIC were extracted for each of the sequenced adenomas. Next, driver genes were prioritised with their previous associations as a colorectal cancer driver by TCGA and Dietlein et al.^16,17^ To compare driver genes and mutation types, genes that were mutated in at least two independent adenomas from either the MBD4-deficient cases or sporadic adenomas were plotted in an oncoprint. In addition, we extracted all somatic mutations present in *APC* from the TCGA Colon Adenocarcinoma (COAD) and Rectum Adenocarcinoma (READ), and from a series of adenomas from patients without germline mutation *APC* mutations.

## RESULTS

### Identification of a colorectal polyposis patient with a biallelic germline MBD4 loss-of-function variant

We performed WGS of constitutional DNA from 35 unrelated patients with at least 10 colonic adenomas before the age of 60, but without pathogenic variants in the known Mendelian CRC genes.^1^ By prioritizing coding germline variants with a high probability of LOF effects we identified a homozygous 4-bp frameshift deletion in *MBD4* (NM_003925.2:c.612_615del; NP_003916.1:p.(Ser205ThrfsTer9)) in a single patient (D:II-1, **Figure 1A**). D:II-1 had no family history of polyposis or CRC and homozygosity analysis did not suggest consanguinity. We found both parents and a sibling to be heterozygous for the *MBD4* frameshift variant **(Figure 1A)**. Re-sequencing of 100 controls from the Mediterranean island from which D:II-1’s family originate did not reveal additional carriers. This variant appears once in 250,800 alleles in gnomAD v2.1.1.

**Figure 1:**
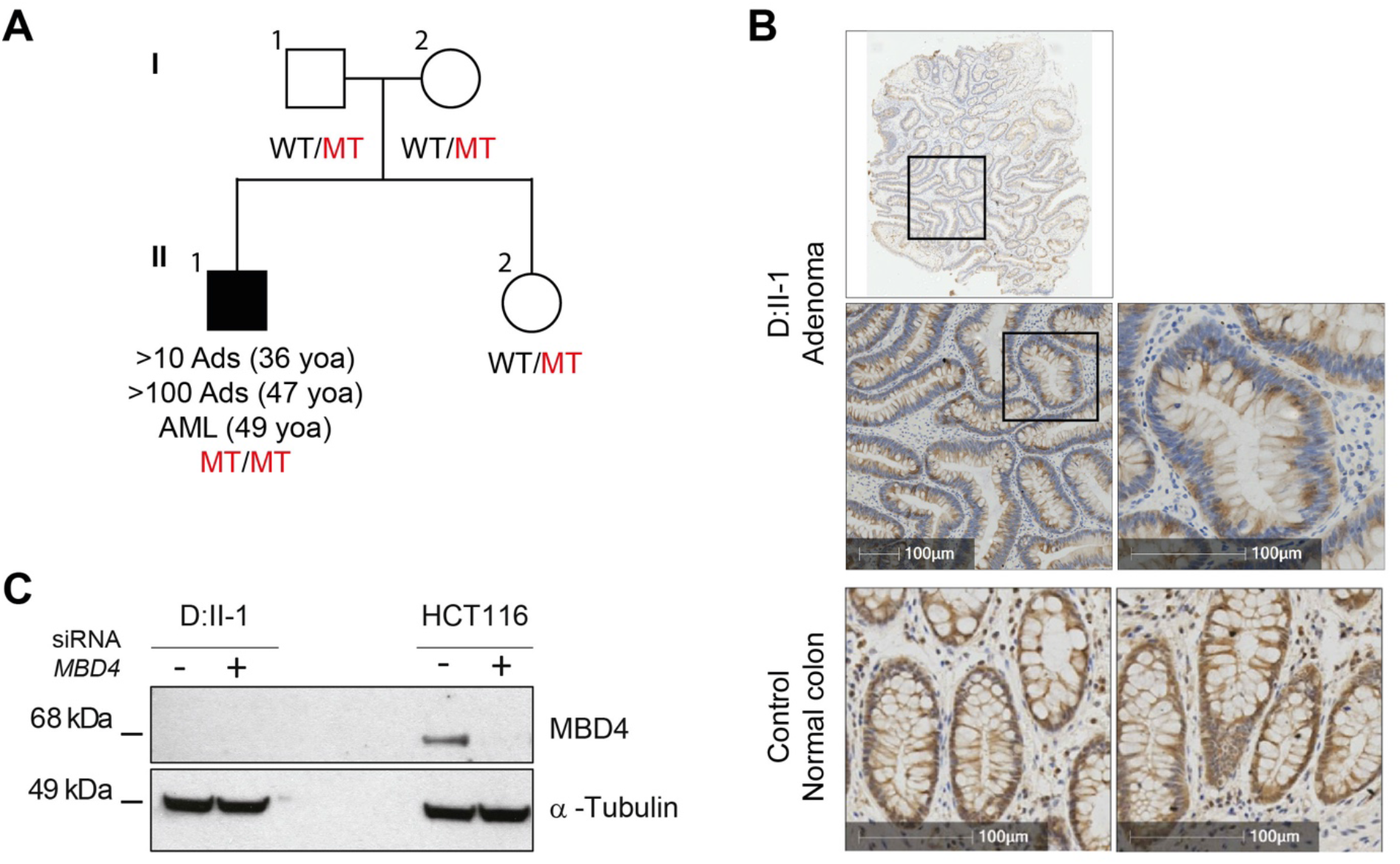
MBD4 deficiency in individual with colorectal adenomas and acute myeloid leukaemia. **A)** Pedigree of Family D with a homozygous *MBD4* loss-of-function variant. For colorectal adenomas, we show the cumulative tumour numbers from age at first presentation and screening colonoscopy to age at last contact. Abbreviations: Ads: adenomas; AML: acute myeloid leukemia; yoa: years of age; MT: mutation; WT: wild-type. **B)** Representative MBD4 IHC of an adenoma from patient D:II-1 (upper panels) and of a normal colon with wild type MBD4 (lower panels) stained with anti-MBD4 antibody. The D:II-1 (upper panels) show a tubular adenoma with low grade dysplasia showing typical nuclear changes (pencil shaped nuclei, crowding and pseudostrastification). **C)** Western blot analysis of MBD4 expression in lymphoblastoid cells from patient D:ll-1 (-) and treated with an siRNA targeting MBD4 (+). The colorectal cancer cell line HCT116 was used a positive control. Two central lanes were left empty. Anti-alpha-Tubulin was used as loading control.

*MBD4* encodes a BER glycosylase that repairs G:T mismatches resulting from the deamination of 5’-methylcytosine (5mC). Immunostaining for MBD4 revealed absence of protein expression in adenomas from D:II-1 **(Figure 1B)**. Furthermore, *MBD4* mRNA was expressed in a lymphoblastoid cell line from D:II-1 **(Supplemental Figure 1A)**, but no protein was detectable **(Figure 1C)**.

D:II-1 presented with rectal bleeding and approximately 60 tubular colorectal adenomas ranging in size from <3mm to 1.5cm diameter at age 36. The largest adenomas were removed and showed evidence of high-grade dysplasia. Following colonoscopy, the patient underwent proctocolectomy at age 47, which identified 50-60 adenomas between the caecum and splenic flexure (mostly in the caecum) and 20-30 rectal adenomas, including some of 1cm-4cm diameter. No gastric or duodenal polyps were identified upon upper gastrointestinal endoscopy screening. Seven months after he underwent proctocolectomy, he presented with pancytopenia and was diagnosed with MDS, which progressed to AML three months later.

### Colorectal phenotype of previously reported AML patients with inherited MBD4 deficiency

During the analysis of D:II-1, in an independent, contemporaneous study, we linked inherited MBD4-deficiency to early-onset acute myeloid leukaemia (AML).^12^ In our prior work, two of the three AML patients with MBD4-deficiency were noted to have developed colonic polyps, whereas this was not evaluated in the third. We have subsequently obtained more comprehensive clinicopathological information on the colorectal phenotypes of those two cases. Patient WEHI-2 (previously WEHI-AML-2) developed a cumulative 17 colorectal polyps over a period of 22 years, with the first polyp excised at the age of 18 years. Histological assessment classified all available polyps (*n=*12) as tubular adenomas with mild-to-moderate dysplasia, with the majority (*n=*7) found in the rectum. A moderately differentiated adenocarcinoma was found in the ascending colon at age 40 and she underwent a right hemicolectomy. EMC-1 (previously EMC-AML-1) developed multiple colonic polyps and underwent a hemicolectomy at age 31 years, although no polyp counts were reported and tissue blocks were unavailable for histological re-assessment.

### Somatic mutation burden, spectra and signatures in MBD4-deficient colorectal adenomas

To investigate the somatic mutation burden and spectrum, we performed WES on nine fresh frozen and two FFPE adenomas from D:II-1, and eight FFPE adenomas from WEHI-2. Some adenomas were micro-dissected to obtain multiple regions of the polyps and normal bowel **(Figure 2A-B; Supplemental Figure 1B-C)**. Almost all somatic mutations in both MBD4-deficient cases were single nucleotide substitutions (SNVs), with a very small proportion of small indels. The fresh frozen adenomas of D:II-1 exhibited an elevated total somatic mutation burden (median 11.1 mutations/Mb [range 8.5-23.3]) compared to a set of nine sporadic adenomas (median 1.8 mutations/Mb [range 1.0-3.1]) **(Figure 2C, Supplemental Figure 2A and Supplemental Table 2)**.^11^ The mutation burden per adenoma increased for both cases with age **(Figure 2D)**. The vast majority of the mutations were CpG>TpG transitions (>95%), which was significantly higher (*P=*2.9*x*10^−7^) than in sporadic colorectal tumours (33% in nine fresh frozen sporadic adenomas, 14% in 468 fresh frozen CRCs from TCGA; **Figure 2E)**.

**Figure 2:**
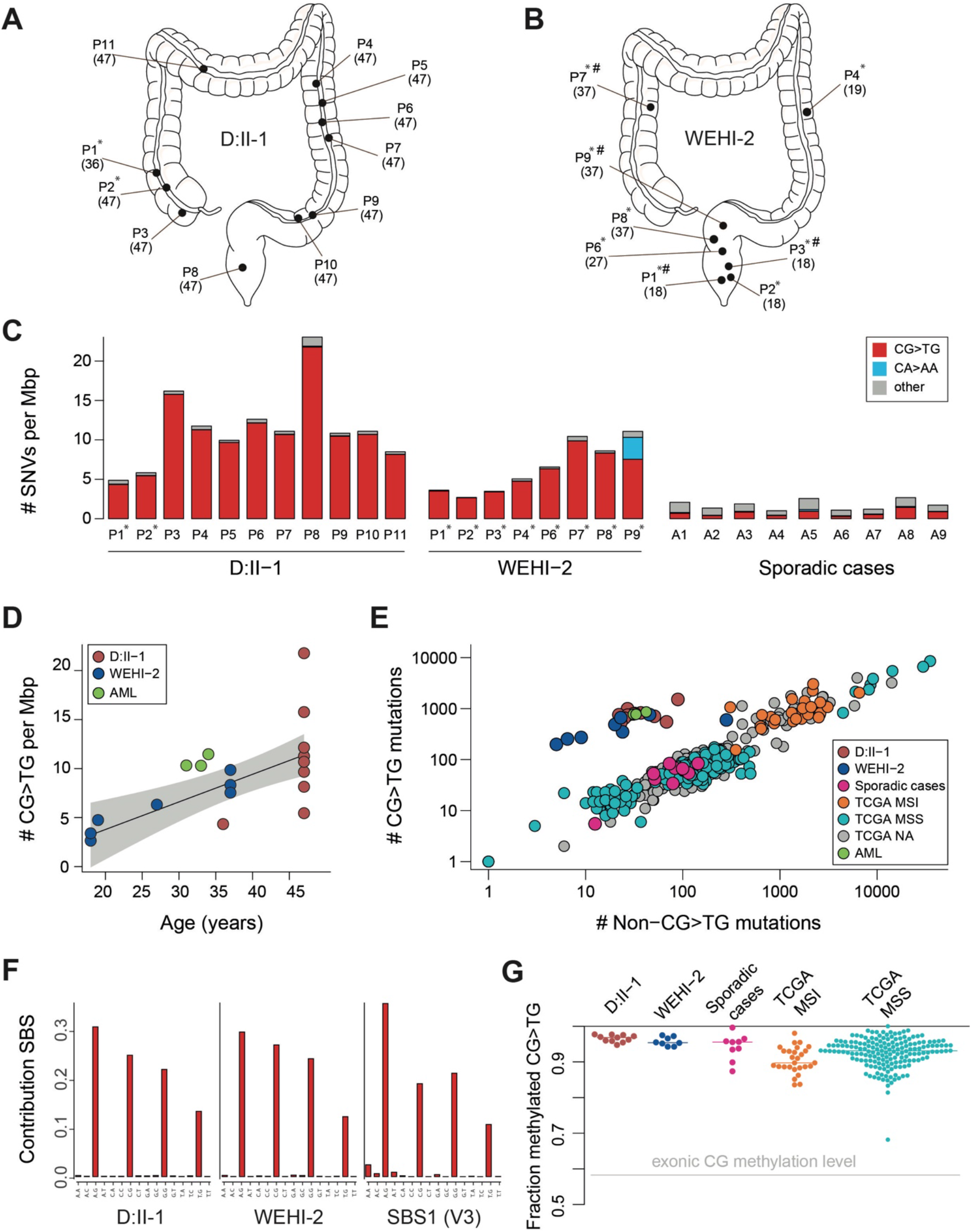
Somatic mutation burden and analysis of polyps of MBD4 deficient cases. **A-B**) Polyps were removed from different parts of the large intestine and rectum from the two MBD4-deficient cases, with age in years specified in brackets. Polyps that were formalin-fixed and paraffin-embedded (FFPE) are indicated with an asterisk (*). Sequencing was performed from multiple regions for some polyps, indicated with a hashtag (#) (H&E shown in Supplemental Figure 1). **C)** Somatic mutation rate for each polyp, FFPE samples indicated with asterisks. The colour of the bars represents mutations in different sequence contexts, with red showing CG>TG mutations, blue showing CA>AA mutations (primarily detected in WEHI-2 P9), and grey representing other base contexts. The median value is presented for samples that had multi-region sequencing. **D)** The number of somatic CG>TG mutations detected in WES data is plotted as function of age. The linear fit is shown, together with an error estimate.**E)** We assessed the contribution of deamination of 5mC to MBD4-deficient samples by comparing the number of CG>TG mutations to all other single nucleotide mutations. The plot compares MBD4 deficient polyps and AMLs ^12^ to sporadic polyps, and to colon and rectal cancers from TCGA.^16^ MSI: microsatellite instability; MSI-H: MSI-high; MSS: microsatellite stable; “MSS” includes both MSS and MSI-low samples; TCGA NA: no MSI data available. **F)** Extracted *de novo* signature SBS1^MBD4^ C>T panel from all polyps from D:II-1(left) and polyps P1-P8 of WEHI-2 (middle), and the C>T panel from COSMIC SBS1-v3 (right). **G)** Fraction of mutated CpG sites that are methylated in normal sigmoid colon (beta value*>*0.5 in WGBS data from the Roadmap Epigenomics Consortium). Each point summarises WES results from a sample and includes all sites with sufficient coverage in WGBS (*n=*177-1507 CG>TG mutations) and the median value is shown with a horizontal line. The grey line shows the fraction of methylated CG sites across all exons.

We performed *de novo* signature analysis on the fresh-frozen adenomas from D:II-1 and separately from the FFPE adenomas from WEHI-2 (excluding P9, see below). The extracted signature profiles were nearly identical to each other (cosine similarity 0.998) and to COSMIC SBS1-v3 (cosine similarity 0.984 and 0.975 for D:II-1 and WEHI-2, respectively; **Figure 2F)**. SBS1-v3 accounted for nearly all somatic mutations detected in D:II-1’s adenomas and most of WEHI-2’s adenomas **(Supplemental Figure 2B)**. The exception was a WEHI-2 rectal polyp (P9) measuring 2cm diameter. All four physically distinct regions of this adenoma had a lower percentage of CpG>TpG transitions (55-77%) than the other adenomas and a substantial proportion of mutations were CA>AA transversions **(Figure 2C)**. Using CpG>TpG transitions as a molecular clock, we found that this shift in mutational profile likely coincided with treatment for AML **(Supplemental Figure 3)**.

**Figure 3:**
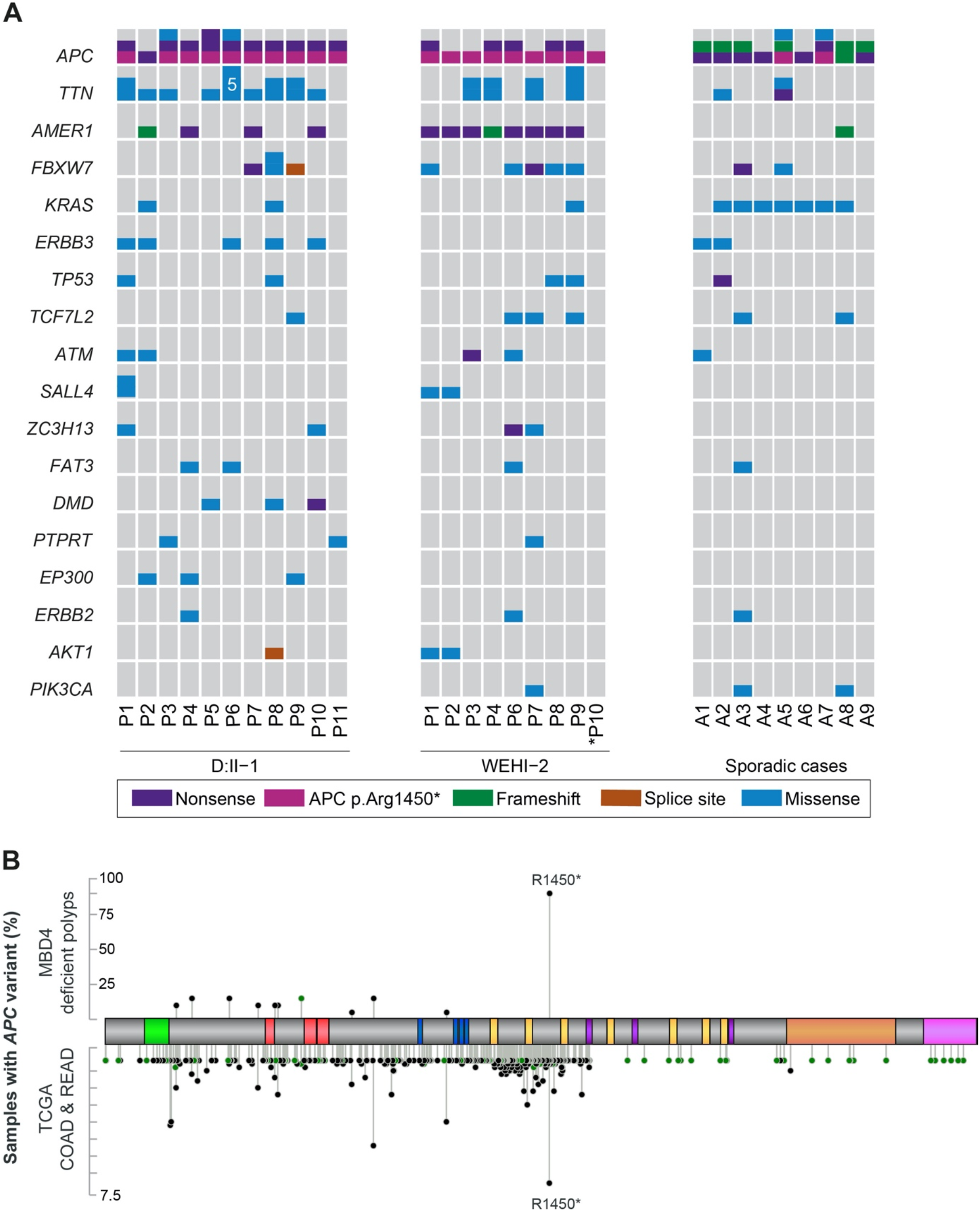
Driver gene mutation analysis of polyps from MBD4-deficient cases. **A)** Oncoprint of driver gene mutation analysis of genes frequently mutated in colorectal cancer. For each polyp the number and type of somatic mutation is shown. * Polyp P10 from WEHI-2 was sequenced using a targeted panel. **B)** Map of the somatic mutations observed in *APC* in all polyps from the two MBD-4 deficient cases and the TCGA COAD and READ datasets. *APC* p.(Arg1450*) is the most frequently observed somatic mutation. Black lollipops indicate protein truncating variants and green lollipops indicate missense variants.

Signature SBS1 is proposed to result from the failure to detect and/or remove G:T mismatches that result from deamination of 5mC. We assessed the methylation status of CpG sites that were mutated in adenomas from D:II-1 and WEHI-2 in whole genome bisulfite sequencing (WGBS) data from normal sigmoid colon generated by the Epigenome Atlas (http://www.roadmapepigenomics.org/). The mutated sites identified in the adenomas from D:II-1 and WEHI-2 were nearly all methylated in normal colon tissue (>96% of sites mutated in both D:II-1 and WEHI-2, compared to 58% across all exonic CpG sites) **(Figure 2G)**.

Next, we compared the mutation profiles of all adenomas of D:II-1 and WEHI-2 to (i) the mutation profiles from the AML samples from EMC-1, WEHI-1 and WEHI-2, (ii) to myeloid progenitors from *Mbd4*^-/-^ mice,^12^ and (iii) to a human HAP1 *MBD4*^ko^ cell line. The somatic mutation profiles and mutation rates were relatively consistent across these diverse samples **(Supplemental Figure 4A-B)**.

### Somatic driver mutations in MBD4-deficient adenomas

AMLs in MBD4-deficient individuals harbour a characteristic set of driver mutations,^12^ and we sought to determine whether this was also true in colorectal adenomas from both D:II-1 (*n=*11) and WEHI-2 (*n=*8). In general, the spectra of driver genes were similar to sporadic CRC and adenomas, although indels were under-represented (most clearly seen for *APC*) **(Supplemental Table 3)**. CRC driver genes with recurrent somatic mutations in our patients’ adenomas included *APC* (*n=*19 adenomas), with a significant enrichment of *APC* p.(Arg1450*), a CpG>TpG transition, compared to sporadic adenomas (Fisher exact, *P=*.0002). In addition, in adenomas from MBD4-deficient cases we observed a significant enrichment of somatic mutations in *TTN* (*n=*13 adenomas; Fisher exact, *P=*.0418) and *AMER1* (*n=*12 adenomas; Fisher exact, *P=*.039) **(Figure 3A)**. Compared to sporadic adenomas, less *KRAS* mutations were observed (3/19 in adenomas from MBD4-deficient cases vs 7/9 in sporadic adenomas; Fisher exact, *P=*.0028). Strikingly, *APC* p.Arg1450* mutations were found in ten out of eleven adenomas from D:II-1 and all of the adenomas sequenced by WES from WEHI-2 **(Figure 3A)**. The preponderance of the *APC* p.Arg1450* mutations unlikely reflects the presence of a shared pre-neoplastic progenitor clone, as the mutation was absent in samples of normal intestine from D:II-1 and WEHI-2 (**Supplemental Table 4**). In sporadic CRCs from TCGA and adenomas from FAP-negative individuals (*n=*31 individuals), p.(Arg1450*) is the most frequent somatic *APC* mutation (32/486 and 8/93 tumours, respectively), but is found in a smaller proportion compared to adenomas from the MBD4-deficient cases (*P<*.00001) **(Figure 3B)**. Public WGBS data indicates similarly high level methylation at multiple CpG hotspot sites within *APC*, suggesting that methylation differences do not explain the over-representation of p.(Arg1450*). Overall, 85.4% and 90.7% of CRC driver mutations in adenomas from D:II-1 and WEHI-2, respectively, were CpG>TpG transitions compared to 36.8% of mutations in nine sporadic adenomas. These data further indicate that 5mC deamination accounts for the vast majority of driver mutations contributing to the development of adenomas in MBD4-deficient cases.

*Most patients with both colorectal tumours and AML do not have germline MBD4 variants* Synchronous or metachronous colorectal polyposis and AML comprise a rare clinical phenotype. In total, we had access to 26 cases with a colorectal and an MDS and/or AML phenotype (see materials and methods) and none had biallelic or monoallelic germline LOF *MBD4* variants.

### Germline LOF MBD4 variants in patients with colorectal polyps and/or CRC

We sought to determine how common MBD4 deficiency is in patients with polyposis and/or CRC. Using a targeted screening strategy, we sequenced the coding sequence of *MBD4* in constitutional DNA from 1,490 European patients with unexplained colorectal polyps (with or without CRC), familial and/or young CRC, or CRC in combination with other tumour(s) (details in **Supplemental Table 1)**. In addition, we assessed 87 families, comprising of 198 CRC- and/or early-onset polyp patients for which the germline was sequenced using WGS/WES for LOF variants in *MBD4*. Finally, we also reviewed 283 constitutional genome sequences from patients with multiple bowel polyps from the pilot programme of the 100KGP. No cases with biallelic germline LOF variants in *MBD4* were identified, but we identified five with heterozygous LOF variants **(Supplemental Table 1; Supplemental Figure 5)**: two nonsense variants (p.(Trp479*); Family A), p.(Arg546*); Family C), one frameshift (p.(Ile111Tyrfs*16); Family F), one synonymous variant that results in defective splicing (c.1410A>C; Family E, **Supplemental Figure 6A-C**) and one canonical splice site (c.1562-1G>T; Family B, **Supplemental Figure 6D-F)**. Compared to the frequency of *MBD4* LOF variant alleles in European non-Finnish gnomAD population this is a significant enrichment (5/3,720 vs 48/129,200; OR =3.62 (95% CI 1.12-9.06), Fisher exact, *P=*.016).

Where possible, we investigated whether the heterozygous *MBD4* LOF variants segregated with the polyposis and/or CRC phenotypes in the respective families. Only one variant (p.(Trp479*), Family A) showed complete co-segregation with the polyposis phenotype **(Supplemental Figure 5)**. For the other families the segregation was discordant or could not (fully) be assessed **(Supplemental Figure 5)**.

To further investigate the impact of heterozygous germline LOF *MBD4* variants on the risk of developing CRC, we studied patients in the CRC Domain of the 100KGP (in which recruitment was not formally enriched for any special features). No carriers of biallelic germline LOF *MBD4* variants were identified. Four Caucasian patients with one LOF allele were identified; this frequency (1.64*x*10^−3^, 4/2,438) was comparable to the frequency observed in a control set of phenotypically disease-free, unrelated participants selected from the 100,000 Genomes Rare Diseases project (1.22*x*10^−3^ (21/17,243), indicating no evidence of enrichment in CRC cases (OR =1.34 (95% CI 0.46-3.93), Fisher exact, *P=*0.59). In the UKBiobank, the frequencies of predicted LOF *MBD4* variants for participants with CRC, colorectal polyps, AML, MDS and all other participants were 8.49*x*10^−4^ (2/2,357), 1.94*x*10^−4^ (1/5,116) 4.5*x*10^−3^ (1/221), 9.6*x*10^−3^ (1/104), 8.64*x*10^−4^ (167/193,255) respectively (as of the October 2020 release of WES data). This data supports no enrichment in any of these groups (Fisher exact, *P>*0.05 for all comparisons).

In contrast to our findings for CRC, germline LOF variants in *MBD4* have been linked to a 9-fold (Fisher exact, *P=*2.00*x*10^−5^) higher incidence of uveal melanoma. Recent reports have shown that these uveal melanomas exhibit loss of the wild-type *MBD4* allele in the tumours.^18-20^ We whole-exome sequenced four colorectal tumors from three individuals with a monoallelic LOF *MBD4* variant and accessed the WGS data of the four CRCs with a monoallelic LOF *MBD4* variant from the 100KGP **(Supplemental Table 1)**. No “second hits” were identified in *MBD4* in any of the tumours. Mutational signature analyses were inconsistent across the four tumours from the three monoallelic *MBD4* LOF variant carriers. In two cases (Family A and F) the mutational signatures were comparable to TCGA COAD samples and the sporadic adenomas **(Supplemental Figure 7)**. However, the two tumours from B:II-1 exhibited a higher contribution of SBS1, more comparable to the MBD4-deficient cases (**Supplemental Figure 7**). However, re-sequencing of *MBD4* in the germline using long-read sequencing did not reveal any additional variants, and Western blot and quantitative real-time PCR suggested the wild-type allele was expressed in blood-derived cell lines from B:II-1 (**Supplemental Figure 6D-F**). We also assessed blood cells from heterozygous *Mbd4* knockout mice and found no evidence of an elevated mutation rate **(Supplementary Figure 4A-B)**. Overall, we found no clear evidence to support a large effect of monoallelic *MBD4* LOF variants on the risk of polyposis and/or CRC.

## DISCUSSION

Our data demonstrate that biallelic germline loss-of-function variants in *MDB4* cause a Mendelian recessive neoplasia syndrome comprising the rare phenotype of both AML and adenomatous colorectal polyposis. Previously, we reported multiple polyps in some AML patients with *MBD4* deficiency, but details of the polyps were not available at that time and hence their importance could not be demonstrated. Here, through independent identification of MBD4-deficiency in a patient with multiple adenomas and more detailed phenotypic and molecular analysis of tumours, we show that inherited MBD4 deficiency causes a true adenomatous polyposis, with a colorectal phenotype similar to attenuated FAP and to patients with germline pathogenic variants in *MUTYH, NTHL1, POLE* and *POLD1*. We suggest the name *MBD4*-associated neoplasia syndrome (MANS) for this condition. The absence of functional MBD4 leads to an accumulation of somatic CpG>TpG mutations arising from spontaneous deamination of 5-methylcytosine, resulting in a somatic mutation spectrum very similar to COSMIC signature v3 SBS1. These mutations include characteristic driver mutations that support the development of adenomas and myeloid leukaemia.

Three out of four individuals with MANS thus far identified developed colorectal polyps early in life, and all four developed MDS/AML. We suggest that *MBD4* coding and splice regions are added to the gene panels currently used for the clinical genetic diagnostic testing of either early-onset AML, or CRC/polyp cases or families. In addition, it would be reasonable to test any individual with multiple polyps and MDS/AML. The pathogenic *MBD4* variants reported to date have included truncating and splice site variants, as well as an in-frame deletion affecting the glycosylase domain of MBD4. However, it remains possible that missense and other variants impair the function of MBD4.^20^

For *MBD4*, we estimate that the frequency of biallelic LOF variants in the general population (based on gnomAD and 100KGP) may be as low as one in 1 in 5-6 million individuals. As is typical of very rare cancer syndromes, it is difficult to estimate age-dependent risks of AML and colorectal tumours in MANS with any precision, although adenomatous colorectal polyps may be present in the second decade of life. While subject to revision as information on the features of MANS accrues, we propose two-yearly colonoscopy from age 18-20 or the date of diagnosis, a regimen often used for other polyposis syndromes caused by deficiency of *MUTYH* or *NTHL1*.^21, 22^ Regular colonoscopy may avoid the need for prophylactic or therapeutic colectomy, especially in patients with relatively low adenoma burdens. Upper gastrointestinal endoscopy is warranted in almost all other polyposis syndromes and it may be prudent to survey the stomach and duodenum at the current time, despite the lack of small bowel or gastric lesions reported in our patients. At least one of the AMLs in our cases developed from MDS and we have also observed clonal haematopoiesis in some cases.^12^ We suggest regular follow-up full blood counts for MANS cases if their initial presentation is with adenomatous polyposis. If the individual presents with AML, then we suggest genetic testing for any family member being considered as a haematopoietic stem cell donor, in keeping with current expert recommendations for managing inherited predisposition to myeloid malignancy.^23,24^

Although carriers of heterozygous LOF *MBD4* variants are susceptible to uveal melanoma,^18-20^ our data shows no convincing evidence that such individuals have a large increased risk of developing polyposis and/or CRC. Complete inactivation of MBD4 is observed in uveal melanoma, where wildtype *MBD4* is lost due to monosomy of chromosome 3, whereas in CRC this is not a frequent event.^25^ Prior analysis of tumours from heterozygous *MBD4* cases in TCGA revealed no significant increase in CpG>TpG mutations,^12^ unless the wildtype allele was inactivated, and this was also consistent with results generated in the *Mbd4* knockout mouse. We cannot, however, rule out the possibility that individuals heterozygous for an MBD4 LOF variant have a small increased risk of CRC and/or polyposis, as has been described for heterozygous LOF *MUTYH* variant carriers.^26^ We identified a single heterozygous LOF *MBD4* case with a substantial number of somatic variants that fit with an MBD4 deficiency in the absence of a second hit in *MBD4* (germline or somatic), but it remains possible that the wildtype allele was silenced.^27^ On balance, it seems unlikely that heterozygous *MBD4* LOF carriers would benefit from colorectal surveillance beyond population screening, unless the individual’s phenotype or family history suggests otherwise.

In conclusion, constitutional deficiency of MBD4 causes a rare genetic syndrome that is marked by the development of adenomatous polyposis and AML. Genetic testing for MANS could be implemented readily and rapidly by incorporating *MBD4* into existing gene panels. MBD4 deficiency results in an elevated mutation burden with a mutation spectrum very similar to COSMIC signature SBS1-v3. A high mutational burden is associated with a good prognosis from CRC, and we speculate that MANS CRCs may also respond well to immune checkpoint inhibitors, as has been reported for PD-1 inhibitors in MBD4-deficient uveal melanomas.^18,19^ It is possible that such a strategy could be used to treat neoplasia in MANS.

## Acknowledgements

We thank Dr. Robbert D.A. Weren, Dr. Janet Vos and Eveline Kamping for technical assistance, the Genome Technology Platform for MIP sequencing support, Dr. Christian Gillissen for the use of the annotation pipeline in the Radboudumc, Nijmegen, and Dr. Peggy Manders for the use of samples from the Dutch Parelsnoer Institute Biobank Hereditary Colorectal Cancer. We acknowledge expert technical assistance from the Austin Molecular Laboratory and the Australian Genome Research Facility for providing expert technical assistance with genomic analysis of cancer samples. We thank members of the Colorectal Oncogenomics Group and the participants and staff from the Colon-CFR in particular, Maggie Angelakos, Samantha Fox and Allyson Templeton for their support of this study. Computation and bioinformatics were also provided by Melbourne Bioinformatics on its Peak Computing Facility. Furthermore, we thank the Birmingham Genomics Service at The University of Birmingham, for the generation of the sequencing and methylation array data. Some of the computations described in this paper were performed using the University of Birmingham’s BlueBEAR HPC service, which provides a High Performance Computing service to the University’s research community. We thank all participants of the CORGI study for their collaboration. The author(s) would further like to acknowledge networking support by the Cooperation in Science and Technology Action CA17118, supported by the European Cooperation in Science and Technology. This research was made possible through access to the data and findings generated by the 100,000 Genomes Project. The 100,000 Genomes Project is managed by Genomics England Limited (a wholly owned company of the Department of Health and Social Care). The 100,000 Genomes Project is funded by the National Institute for Health Research and NHS England. The Wellcome Trust, Cancer Research UK and the Medical Research Council have also funded research infrastructure. The 100,000 Genomes Project uses data provided by patients and collected by the National Health Service as part of their care and support. The colon schematics presented in Figure 2 was sourced from Servier SMART Medical Art (https://smart.servier.com/) and modified for us under a CC BY 3.0 license.

## Contributors

Study supervision: CP, IPMT, IJM, RMdV. Analysis and drafting: CP, EC, JEG, SG, CF, IPMT, IJM, RMdV. Data support: EAMJ, HC, LC, EAB, AB, KS, LM, HT, DG, FF, YG, DJA, SAMvdB, MC, MC, CD, JAD, DD, JL, KM, RSvdP, MS, JW, JCT, CT, LV, TvW, CW, CA, GC, WC, DC, AF, AG, BK, BN, DC, TG, RH, NLB, AS, DW, MAJ, RPK, AWR, MJLL, NH, VHK, ADR, IMW, CRP, DDD. Critical revision: all authors. Shared last authors: IPMT, IJM, RMdV.

## Appendix

### ^a^Genomics England Research Consortium

Ambrose, JC^1^, Arumugam, P^1^, Bleda, M^1^, Boardman-Pretty, F^1,2^, Boustred, CR^1^, Brittain, H.^1,^ Caulfield, MJ.^1,2^, Chan, GC^1^, Fowler, T^1^, Giess A^1^, Hamblin, A.^1^, Henderson, S.^1,2^; Hubbard, TJP^1^, Jackson, R^1^, Jones, LJ^1,2^; Kasperaviciute, D^1,2^, Kayikci, M^1^, Kousathanas, A^1^, Lahnstein, L^1^, Leigh, SEA^1^, Leong, IUS^1^, Lopez, FJ^1^, Maleady-Crowe, F^1^, Moutsianas, L^1,2^, Mueller, M^1,2^, Murugaesu, N^1^, Need, AC^1,2^, O‘Donovan P^1^, Odhams, CA^1^, Patch, C^1,2^, Perez-Gil, D^1,^ Pereira, MB.^1^, Pullinger, J^1^, Rahim, T^1^, Rendon, A^1^, Rogers, T^1^, Savage, K^1^, Sawant, K^1^, Scott, RH^1^, Siddiq, A^1^, Sieghart, A^1^, Smith, SC^1^, Sosinsky, A^1,2^, Stuckey, A^1^, Tanguy M^1^, Thomas, ERA^1,2^, Thompson, SR^1^, Tucci, A^1,2^, Walsh, E^1^, Welland, MJ^1^, Williams, E^1^, Witkowska, K^1,2^, Wood, SM^1,2^.^*1*^ *Genomics England, London, UK;* ^*2*^ *William Harvey Research Institute, Queen Mary University of London, London, EC1M 6BQ, UK*.

### ^b^The CORGI Consortium

Dr. Kai Ren Ong (Birmingham Women’s Hospital, Birmingham), Prof. Andrew Beggs (Institute of Cancer and Genomic Sciences, University of Birmingham), Dr. Alan Donaldson (St. Michael’s Hospital, Dr. Ruth Armstrong, Addenbrooke’s NHS Trust, Cambridge), Dr. Carole Brewer (Royal Devon & Exeter Hospital (Heavitree)), Exeter, Prof. Jayantha Arnold (Ealing Hospital, Middlesex), Dr. Munaza Ahmed (Great Ormond Street Hospital, London), Dr. Louise Izatt (Guy’s Hospital, London), Dr. Andrew Latchford (St Mark’s Hospital, Harrow and Division of Surgery and Cancer, Imperial College London), Dr. Dorothy Halliday (Nuffield Orthopaedic Hospital, Oxford), Peter Risby (The Oxford Genomic Medicine Centre, Oxford), Dr. Paul Brennan (The James Cook University Hospital, Middlesbrough), Dr. Alison Kraus (Chapel Allerton Hospital, Leeds), Dr. Julian Barwell (Leicester Royal Infirmary, Leicester), Dr. Lynn Greenhalgh (Liverpool Women’s Hospital), Prof. D. Gareth Evans (University of Manchester, Manchester), Kate Green (University of Manchester, Dr. Timothy Simmons (Institute of Genetic Medicine, International Centre for Life, Newcastle upon Tyne), Dr. Rachel Harrison (Nottingham University Hospitals, Nottingham), Prof. Ragunath (Queen’s Medical Centre Campus, Nottingham), Prof. Brian Davidson (Royal Free Hampstead NHS Trust, University Dept. of Liver Medicine & Transplantation, London), Dr. Zoe Kemp (The Royal Marsden, Sutton), Dr. Helen Hanson (St George’s University, London), Dr. Katie Snape (St George’s University, London), Prof. Anneke Lucassen (Princess Anne Hospital, Southampton), Dr. Kevin J Monahan (West Middlesex University Hospital, Middlesex), Prof. Patrick Morrison (City Hospital Campus, Belfast).

### ^c^WGS500 Consortium

Peter Donnelly^1^, John Bell^2^, David Bentley^3^, Gil McVean^1^, Peter Ratcliffe^1^, Jenny Taylor^1,4^, Andrew Wilkie^4,5^, John Broxholme^1^, David Buck^1^, Jean-Baptiste Cazier^6^, Richard Cornall^1^, Lorna Gregory^1^, Julian Knight^1^, Gerton Lunter^1^, Ian Tomlinson^7^, Andrew Wilkie^4, 5^, Christopher Allan^1^, Moustafa Attar^1^, Angie Green^1^, Lorna Gregory^1^, Sean Humphray^3,^ Zoya Kingsbury^3^, Sarah Lamble^1^, Lorne Lonie^1^, Alistair Pagnamenta^1^, Paolo Piazza^15^, Amy Trebes^1^;, John Broxholme^1^, Richard Copley^8^, Simon Fiddy^9^, Russell Grocock^3^, Edouard Hatton^1^, Chris Holmes^1^, Linda Hughes^1^, Peter Humburg^1^, Alexander Kanapin^10^, Stefano Lise^11^, Hilary Martin^12^, Lisa Murray^3^, Davis McCarthy^13^, Andy Rimmer^14^, Natasha Sahgal^1^, Ben Wright^1^, Chris Yau^16. *1*^*The Wellcome Trust Centre for Human Genetics, Roosevelt Drive, Oxford, OX3 7BN, UK;* ^*2*^*Office of the Regius Professor of Medicine, Richard Doll Building, Roosevelt Drive, Oxford, OX3 7LF, UK;* ^*3*^*Illumina Cambridge Ltd*., *Chesterford Research Park, Little Chesterford, Essex, CB10 1XL, UK;* ^*4*^*NIHR Oxford Biomedical Research Centre, Oxford, UK;* ^*5*^*Weatherall Inst of Molecular Medicine, University of Oxford; John Radcliffe Hospital,Headington, Oxford OX3 9DS, UK;* ^*6*^*Centre for Computational Biology, Haworth Building, University of Birmingham, Edgbaston, Birmingham B15 2TT, UK;* ^*7*^*Edinburgh Cancer Research Centre, IGMM, University of Edinburgh, Crewe Road, Edinburgh EH4 2XR, UK;* ^*8*^*UPMC Paris 6, CNRS UMR 7009, Villefranche-sur-Mer;* ^*9*^*Oxford Nanopore Technology, Edmund Cartwright House, 4 Robert Robinson Avenue, Oxford Science Park, Oxford, OX4 4GA, UK;* ^*10*^*CRUK Oxford Centre, Department of Oncology, Old Road Campus Research Building, Roosevelt Drive Oxford, OX3 7DQ, UK;* ^*11*^*Centre for Evolution and Cancer, The Institute of Cancer Research, Brookes Lawley Building, 15 Cotswold Road, Sutton, SM2 5NG, UK;* ^*12*^*Sanger Institute, Wellcome Genome Campus, Hinxton, Cambridge, CB10 1SA, UK;* ^*13*^*EMBL-EBI, Wellcome Genome Campus, Hinxton, Cambridgeshire, CB10 1SD, UK;* ^*14*^*Genomics plc, King Charles House, Park End Street, Oxford, OX1 1JD, UK;* ^*15*^*Imperial College London, Commonwealth Building, Hammersmith Campus, Du Cane Road, London, W12 0NN, UK;* ^*16*^ *Division of Informatics, Imaging & Data Sciences, University of Manchester, UK*

## SUPPLEMENTAL FILES

### Supplementary Methods

**Supplemental Figure 1:** MBD4 expression in lymphoblastoid cells and histology of polyps from MBD4 deficient cases

**Supplemental Figure 2:** Somatic burden and mutational signature refitting of adenomas of MBD4 deficient cases.

**Supplemental Figure 3:** Clonal evolution of adenomas in WEHI-2.

**Supplemental Figure 4:** Mutation profiles and cosine similarity of observed mutations in various samples.

**Supplemental Figure 5:** Pedigrees of heterozygous *MBD4* loss-of-function variant carriers and co-segregation analyses.

**Supplemental Figure 6:** Analyses of MBD4 heterozygous splice site variants.

**Supplemental Figure 7:** Mutational signature refitting analyses of tumours from heterozygous MBD4 loss-of-function variant carriers.

**Supplemental Table 1:** Patient cohort inclusion and results of the molecular inversion probe screening.

**Supplemental Table 2:** Details on the tumours subjected to sequencing.

**Supplemental Table 3:** Somatic driver mutations in genes associated with colorectal cancer development in MBD4-deficient and sporadic polyps.

**Supplemental Table 4:** Validation of somatic driver mutations in by panel sequencing in polyps and normal tissues from WEHI-2.

