## Supplemental material for "Germline loss-of-function variants in the base-excision repair gene *MBD4* cause a Mendelian recessive syndrome of adenomatous colorectal polyposis and acute myeloid leukaemia"

32. Department of Medicine, Melbourne Medical School, Faculty of Medicine, Dentistry and Health Sciences, University of Melbourne, Melbourne, Australia
33. Fundación Pública Galega de Medicina Xenómica SERGAS, Grupo de Medicina Xenómica-USC, Instituto de Investigación Sanitaria de Santiago (IDIS), Centro de Investigación Biomédica en Red de Enfermedades Raras (CIBERER), Santiago de Compostela, Galicia, Spain
34. University of Melbourne, Department of Medical Biology, 1G Royal Parade VIC 3052 Australia
[a,b, c] See appendix
* *These authors contributed equally;* [#] *These authors jointly supervised this project*

#### Supplementary Methods:

Whole-genome sequencing for germline variants

Patients with multiple colorectal adenomas were recruited as part of the UK CORGI study. As part of the Oxford-Illumina WGS500 project, we performed whole-genome sequencing of constitutional DNA from 35 probands with at least 10 colonic adenomas before age 60.[1] The Illumina HiSeq platform was used and a median of ~40X coverage achieved. Read alignment and variant calling were performed using BWA, Stampy and Platypus as described.[1] Samples were additionally joint called using GATKv3 and annotated using ANNOVAR.[1] We extracted all variants predicted to result in protein truncation (nonsense, frameshift, or splice site variants) and prioritized for homozygous and compound heterozygous variants with a low frequency (MAF < 0.01) in the general population.

Molecular inversion probe sequencing

All participants in the MIP screen provided written informed consent. This study was approved by the local medical ethics committee (CMO light; study number 2015/2172 of the Radboudumc Nijmegen). Leukocyte-derived DNA was used for targeted resequencing of *MBD4* (NM_003925.2) using 32 Molecular Inversion Probes (MIPs), covering all coding regions and intron-exon boundaries, were designed according to the previously published methodology.[2, 3] After targeted capture, samples were sequenced on a NextSeq500 (Illumina) system. Reads were mapped using BWA and variants called using GATK's UnifiedGenotyper. After variant calling, all variants with an at least 40-fold absolute coverage, ≥20 variant reads, ≥25% variant reads and ≥8.000 quality by depth scores were selected for further analyses. Loss-of-function (LOF) variants in *MBD4* (see below) identified using MIP-sequencing with a quality by depth score of 8.000-11.000 were validated using Sanger sequencing.

MBD4 loss-of-function variant prediction and -selection

All *MBD4* variants located in an exon region, canonical splice site (positions +1, +2 and −2, -1), and coding or noncoding splice site region (3' splice site -12 till +2 and 5' splice site -3 till +6) were included for further analyses. Furthermore, only variants with an allele frequency <1% in an in-house database of 12,244 germline exomes that have been sequenced at Genome

Diagnostics Nijmegen (https://order.radboudumc.nl/en/genetics) and <2% in ExAC and gnomAD were included. To select variants of pathogenic potential, we selected all frameshift and nonsense variants, and missense variants with a PhyloP score ≥3 and a CADD_PHRED score ≥15. For variants with a predicted splicing effect of more than 20% by SpliceAI additional in silico splice site predictions were obtained using MaxEntScan, NNSPLICE, and Human Splicing Finder (Alamut Visual 2.13). Splice site losses were included when 1) the variant splice score was less than 50% of the scoring range for at least two algorithms and 2) the difference between wildtype and variant splice score was more than 20% of the scoring range in at least three algorithms. Splice site gains were included when 1) the variant splice score was above 75% of the scoring range for at least two algorithms and 2) the difference between the gained splice site and the nearest splice site was more than 2% of the scoring range in at least two algorithms.

Generation of a lymphoblastoid cell line from patient D:II-1

Peripheral blood lymphocytes (PBLs) were isolated using Ficoll-Paque PLUS (eppendorf) following manufacturers instructions from a fresh blood sample, collected in sodium heparin tubes from patient D:II-1. A lymphoblastoid line was generated by Epstein Barr virus transformation by the Culture collections team, Public Health England.

RNA and protein analysis

Taqman expression probes HS01023548 and HS00187498 were used to quantify *MBD4 m*RNA extracted from a lymphoblastoid cell line from patient D:II-1 (further details available upon request). Protein lysates from cells were analysed by western blotting using anti-MBD4 antibody ab224809 (Abcam). HCT116, HAP1 and D:II-1 lymphoblastoid cells were resuspended in RIPA buffer (Thermo Fisher Scientific). Total lysates were quantified with Pierce BCA Protein Assay kit (Thermo Fisher Scientific) according to manufacturer's instructions. 20 µg of protein lysate were loaded on a 4-20% gradient gel (Thermo Fisher Scientific) or NuPage 4-12% Bis-Tris Gels (Invitrogen). After transfer with iBlot2 dry Blotting System (Thermo Fisher Scientific) and blocking membranes were blotted for anti-MBD4 (abcam, diluted 1:1000) and anti-α-tubulin (Sigma, 1 diluted 1:5000 or Abcam, diluted 1:500) as a loading control. Membranes were exposed to hyperfilm ECL (GE Healthcare) and developed using a X-Ray Film Processor (Konica) or scanned on the Odyssey Infrared Imaging System (Li-COR).

MBD4 immunohistochemistry

Formalin-fixed paraffin-embedded (FFPE) specimens of colorectal adenomas were analysed by immunohistochemistry using the same anti-MBD4 antibody. Samples sections (5 µm) were deparaffinized with xylene and rehydrated. Sections were incubated with 6% $H_2O_2$ for 20 min at room temperature to block endogenous peroxidase activity. Antigen retrieval was carried out by incubating the slides in citrate buffer (pH 6.0) at 95°C for 10 min. Sections were blocked with goat serum for 30 min at room temperature. Primary antibodies were diluted in 1% goat serum/0.1% BSA/PBS. Sections were incubated with MBD4 primary antibody (ab224809, 1:50) overnight at 4°C. Sections were washed with Tris-buffered saline with 0.1% Tween 20 (TBST) and incubated with secondary antibody anti rabbit (PK-6100) for 30 min. Tertiary (ABC Biotinylated) was kept for 30 min. Staining was visualized using a HRP/DAB detection system Dako. Control IHC experiments (data not shown) were performed without primary antibody. All sections were counterstained with Gill's haematoxylin and mounted for digital slide scanning using a Zeiss ActioScan Z1.

*In vitro* splicing assay

To determine if the synonymous *MBD4* variant c.1410A>C has an effect on splicing we used a minigene assay as described by Sangermano *et al*.[4] In brief, *MBD4* exon 2-7 (GRCh37; chr3:129151044-129157184) was amplified from gDNA of E:II-1and cloned into a minigene expression vector. Both wildtype and mutant minigenes were transfected into HEK293T cells and HAP1 *MBD4*$^{KO}$ cells. After 36h, total RNA was extracted and cDNA was generated as described previously.[5] Ultimately, the splicing effects of both minigenes were measured by reverse transcriptase (RT) PCR on cDNA isolated from the transfected cells (**Supplemental Figure 6**).

Genomic qPCR

Analyses of genomic copy number variants were performed using quantitative PCR (qPCR) using probes that target all exons of *MBD4* on a 7500 Fast Real-Time PCR system (Applied Biosystems) as described previously.[5]

Long-read sequencing of *MBD4*

The full *MBD4* locus was amplified by long-range PCR with primers targeting the complete *MBD4* locus and at least 2kb of the up- and downstream flanking regions using LongAmp (New England Biolabs). The resulting PCR product was purified using 1.5x AMPure PB beads (Pacific BioSciences) and subjected to long-read SMRT sequencing using the 'Procedure & Checklist – Preparing SMRTbell™ Libraries using PacBio® Barcoded Adapters for Multiplex SMRT® Sequencing' protocol (Pacific BioSciences) on the PacBio Sequel I System. Circular consensus sequence (CCS) analyses and mapping to the reference genome (hg19) was performed using SMRT link v6.1 (Pacific BioSciences) with standard parameters with the exception of applying a 'minimal predicted accuracy' of 0.99 and the 'placeGapConsistently' parameter in the advanced settings. Subsequently, bam files were further analyzed using IGV and SeqNext (JSI medical systems; version 4.4.0, build 509).

Whole-exome sequencing of adenomas

WEHI-2 (previously reported as WEHI-AML-2) consented to the use of their clinical information and tissues for research in accordance with the Declaration of Helsinki. The project was approved by human research ethics committees from the Walter & Eliza Hall Institute of Medical Research (WEHI) and Melbourne Health (MH) (WEHI HREC project 13/01, MH HREC project 2012.274). DNA was extracted from nine fresh frozen adenomas, two formalin fixed paraffin embedded (FFPE) adenomas and three fresh frozen macroscopically normal bowel tissues from D:II-1, nine independent FFPE adenomas from WEHI-2, and FFPE tumours from the heterozygous cases and a wild-type colorectal cancer (CRC) **(Supplemental Figure 1B and 1C; Supplemental Table 2)**. Exome library preparations were performed according to the manufacturer using either the i) Agilent SureSelectXT Human All Exon V6 (Agilent Technologies), ii) Agilent SureSelect Clinical Research Exome version 2 (Agilent Technologies), iii) Agilent SureSelect XT Low Input Human Whole Exome V6 (Agilent Technologies) or iv) Illumina TruSeq exome (Illumina) enrichment kit in combination with sequencing on a NextSeq500 (Illumina) or NovaSeq 6000 (Illumina) **(Supplementary Table 2)**.

WES sequencing reads from D:II-1 were aligned with BWA and stampy v1.0.28. Duplicates were marked using picard 1.9.2. Clonal tracking was performed with default superFreq (v1.3.2)[6] using preliminary variants from VarScan v2.3[7] with the options --strand-filter 0, --p-value 0.05 and --min-var-freq 0.02. For the FFPE samples >700,000 variants were called per sample using these settings compared to ~100,000 in the fresh frozen samples. Varscan VCFs from FFPE samples were additionally filtered to remove variants with <10% VAF and <4 reads supporting the variant allele. Somatic variants were called using Mutect2 (GATK version 4.1.0.0). Normal bowel samples from the caecum and transverse colon of patient D:II-1 were used as matched normal, a panel of 6 normal colon samples from other patients without CRC or polyps were used as a panel of normals reference to assist with filtering platform artefacts. "af-only-gnomad.raw.sites.b37.vcf" was provided as an additional germline reference.

WES sequencing reads from WEHI-2 were aligned to hg19 with BWA.[8] Variant calling, CNA calling and clonal tracking was done with default superFreq (v1.3.2)[6] using preliminary variants from VarScan (v2.3)[7] with the options --strand-filter 0, --p-value 0.01 and --min-var-freq 0.05. To limit artefacts, the variants were further filtered based on clonal assignment.

WES sequencing reads from the heterozygous cases were aligned with BWA. Variants were called with Mutect2 (GATK version 4.1.0.0), either with a matched normal exome derived from healthy tissue **(Supplementary Table 2)**. A panel of 100 germline samples from a control population without CRC or polyps was used as a reference to assist with filtering platform artefacts. Somatic variants were selected as described for D:II-1, with the minor modification of excluding variants ≤2 reported in ExAC.

WES methods for the sporadic adenoma samples was previously described.[9] Somatic variants were called from the BAMs for each region, which were previously aligned to hg19, using Mutect2, following the same strategy as described for D:II-1. Each adenoma sample had a matched normal. In all analyses the union of mutations called in any of the regions from the same adenoma were combined.

WES data from WEHI-2 and D:II-1 will be made available through EGA (EGAS00001004842 and EGAS00001005063). Data is available for ethically approved cancer research upon completion of a data transfer agreement, which includes restrictions on the disclosure of germline variants. Somatic variants in selected driver genes are available in **Supplementary Table 3**. Somatic variant calls are available upon request.

Comparison to TCGA CRC

To compare to the The Cancer Genome Atlas (TCGA) CRC data, we downloaded variant calls from SomaticSniper, VarScan2, MuTect2, and MuSE through the National Cancer Institute Genomic Data Commons. Variants were retained if the variant allele frequency was greater than 20% with at least 20 read depth, and if it was identified by at least 3 of the 4 callers. Mismatch repair status was available for a subset of samples,[10] which we used to classify patients as microsatellite stable (MSS) or unstable (MSI).

CRISPR/Cas9 generated *MBD4* knockout cells

HAP1 cells were maintained in Iscove's Modified Dulbecco's Medium (IMDM; GIBCO), containing 10% fetal calf serum (FCS), 1% glutamine, and 1% penicillin/streptomycin. Single guide RNAs (sgRNA) targeting the glycosylase domain of *MBD4* were designed using CHOPCHOP,[11] cloned into the Cas9 expression vector PX459 v2.0 (Addgene plasmid #62988) and HAP1 cells were transfected according to the method described by Ran *et al.* with minor modifications.[12] After puromycin selection single cell clones were derived using a FACS sorting. Effective knockout of *MBD4* was determined based on Sanger sequencing of the target regions, mRNA expression and by Western blot. Full *MBD4* knockout (*MBD4*$^{KO}$) HAP1 single cell clones were cultured for 142 days, followed by another single cell sort. Subclones were expanded for 14 days and gDNA was isolated from the parental HAP1 clone and *MBD4*$^{KO}$ subclones. Two WT and *MBD4*$^{KO}$ clones were whole-genome sequenced using the TruSeq DNA PCR-Free library kit (Illumina) and sequenced on a NovaSeq 6000 System (Illumina). Reads were mapped using BWA and for all samples at least 90% of the genome was covered at 20X. Variant calling and mutational signature analysis was performed as described above. As modification to the analysis the average 96-profile of the HAP1 WT clones was extracted from the *MBD4*$^{KO}$ clone to be left with the 96-profile specific to the *MBD4* knockout.

Assessment of mutation rate in a mouse model of MBD4 deficiency

Whole genome sequencing was performed on individual mouse haematopoietic progenitor colonies as previously described.[13] In brief, mouse bone marrow cells were cultured in semi solid agar. Each culture contained 10,000 bone marrow cells, suspended in Dulbecco modified Eagle medium with 20% bovine calf serum and 0.3% agar, with 100ng murine stem cell factor, 10ng murine IL-3 and 2 IU erythropoietin. Cells were incubated for 11 days at 37°C in a humidified atmosphere with 10% $CO_2$. DNA was extracted from individual colonies using QIAamp DNA Micro Kit (Qiagen), amplified using TruePrime WGA Kit (SYGNIS) and purified using QIAamp DNA Mini Kit (Qiagen). Whole genome sequencing was performed on the NovaSeq 6000 (150bp paired end reads, Illumina)**.** The mouse sequencing data was aligned to the mouse genome (mm10) using bwa-mem. WGS was also performed on the original bone marrow DNA and used to identify variants unique to the individual colony. Results from the wildtype and knockout colonies were reported previously and were deposited at SRA (Accession: PRJNA419992).[13]

UK 100,000 genomes samples

Whole genome sequencing data from constitutional DNA was available for 243 multiple (>20) colorectal adenoma cases and 2,646 colorectal cancer cases. Genomes were sequenced on the Illumina HiSeq X platform and reads aligned to the human genome (GRCh38) using the Illumina iSAAC aligner 03.16.02.1.[14] Mutations were called using Starling.[14] Genome VCF files were joint genotyped using gvcfgenotyper (https://github.com/Illumina/gvcfgenotyper) and annotated using VEP. All PASS variants mapping to *MBD4* were extracted. Variants mapping to the 10bp homopolymer tract (chr3:129,436,705-129,436,714) were excluded as likely false positive calls. IBD (KING[15]) analyses were performed on the CRC patient genomes and 21,564 selected rare disease genomes. One individual in each pair with a pi hat value >0.25 was removed. Principal components analysis (PLINK[16]) of the unrelated individuals, together with

HapMap samples was performed to identify genetic based ethnicity. This set of rare disease genomes comprised healthy relatives of probands with syndromes not associated with an increased cancer predisposition. Germline *MBD4* mutation frequencies were derived from unrelated cases (multiple adenomas or CRC) and rare disease controls within the main Caucasian centroid from the first two PCs. Somatic mutation calls generated using Strelka and curated by the Colorectal cancer domain of the 100,000 genome project were interrogated to search for second hits in *MBD4* in the four colorectal patients with a single LOF germline *MBD4* variant.

**LEGENDS TO SUPPLEMENTAL FIGURES:**

**Supplemental Figure 1: MBD4 expression in lymphoblastoid cells and histology of polyps from MBD4 deficient cases. A)** RNA expression analysis showed stable expression of MBD4 as determined using two Taqman probes targeting MBD4 RNA (probe1=hS01023548; probe 2=HS00187498). Ct averages were 29.1 and 29.3 for HCT116 and D:II-1 respectively, using probe_1 and 26.7 and 26.4 for HCT116 and D:II-1 respectively, using probe_2. Bars are plotted as average of triplicate experiments with standard deviation. **B)** Haematoxylin and eosin staining of polyps from D:II-1 with age in brackets. **C)** Haematoxylin and eosin staining and immunohistochemistry in polyps from WEHI-2. Rows 1 & 2: Haematoxylin and eosin (H&E) staining of sections from 10 polyps excised from WEHI-2, with age in years in brackets. Polyps P3, P7, P9 and P10 had multi-regions independently sequenced these regions are indicated by circles. Size bars represent 2000uM (P1 to P6) or 5000um (P7 to P10). Row 3: Immunohistochemistry staining of WEHI-2 P10 showing proficiency in mismatch repair proteins (MLH1, MSH2, MSH6 and PMS2). An H&E slide is shown for comparison. Size bars represent 500uM.

**Supplemental Figure 2: Somatic burden and mutational signature refitting of polyps of MBD4 deficient cases. A)** Fractions of single nucleotide mutations for each polyp. The colour of the bars represents mutations in different sequence contexts, with red showing CG>TG mutations, blue showing CA>AA mutations (primarily detected in WEHI-2 P9), and grey representing other base contexts. Polyps that were formalin-fixed and paraffin-embedded (FFPE) are indicated with an asterisk (*). The median value is presented for samples that had multi-region sequencing. **B)** The estimated relative contribution of all known COSMIC-v3 mutational signatures to mutations in whole-exome sequenced adenomas from two MBD4-deficient cases and nine adenomas from sporadic origin. For WEHI-2 and the sporadic cases for some polyps multi-regions were sequenced. For WEHI-2 multi-region sequenced polyps are indicated by P1A or P1B, ect., and for the sporadic cases this is indicated by by A1.1 and

A1.2, ect. (see also Supplemental Table 2). Signatures with a contribution of less than 5% were merged into “other”.

**Supplemental Figure 3: Clonal evolution of polyps in WEHI-2.** These trees represents the development of clones within each polyp, with vertical bars or branches representing subclones. Note that P1 and P2 share a common precursor; labels are placed adjacent to the dominant clone in each polyp. The x-axis shows the number of somatic CG>TG mutations in each clone. Timing for key driver mutations is shown with earlier mutations on the left and the colour reflecting the type of mutation, either CG>TG (red), CA>AA (blue), copy number (green) or other (grey).

**Supplemental Figure 4: Mutation profiles and cosine similarity of observed mutations in various samples. A)** Combined mutation profiles in the 96-mutation spectrum plot for each of the samples indicated. **B)** Cosine similarity scores indicate the closeness of the mutation profile of D:II-1 and WEHI-2 with the various mutations profiles observed in the other sequenced samples.

**Supplemental Figure 5: Pedigrees of heterozygous *MBD4* loss-of-function variant carriers and co-segregation analyses.** Numbers in brackets indicate ages at which various adenomas were diagnosed. CRC: colorectal cancer; TA: tubulovillous adenomas; GC: gastric cancer; KC: kidney cancer; PrC: prostate cancer; HP: hyperplastic polyps; SA: serrated adenoma; TVP: tubulovillous polyp; AC: adenocarcinoma; A: adenoma.

**Supplemental Figure 6: Analyses of *MBD4* heterozygous splice site variants. A)** Splicing predictions of Alamut Visual 2.10 for the c.1410A>C variant of patient E:II-1. Shown are the scores for SSFL: SpliceSiteFinder-like, HSF: Human Splicing Finder, SpliceAI (Jaganathan et al., 2019), and ESEfinder predictions. **B)** Minigene splicing assay results for the synonymous *MBD4* variant c.1410A>C. Displayed is the agarose gel of the reverse transcriptase PCR product of primers amplifying *MBD4* exons 4 to 7. cDNA input was derived from non-transfected (NT) cells, transfected cells with wildtype (WT) minigene, and transfected cells with mutant (MT) minigene in both HEK293T and HAP1 MBD4KO cells. **C)** The size of the agarose bands corresponds to the predicted cDNA length for the depicted primer pair, which amplifies exons 4 to 7 of MBD4, for the wildtype and mutant allele. **D)** The relative MBD4 mRNA expression of a lymphoblastoid cell line (LCL) derived from patient B:II-1 (LCL B:II-1) compared to four control LCLs. **E)** A representative image of the MBD4 protein expression of LCL B:II-1 and four control LCLs by Western blot. **F)** The relative MBD4 protein expression (n=4) of LCL

B:II-1 compared to four control LCLs. Bars are plotted as average of triplicate experiments with standard deviation.

**Supplemental Figure 7: Mutational signature refitting analyses of adenomas from heterozygous *MBD4* loss-of-function variant carriers.** The estimated relative contribution of all known COSMIC-V3 mutational signatures to mutations in whole-exome sequenced adenomas from wild-type and heterozygous *MBD4* loss-of-function variant carriers from pedigrees in Supplemental Figure 5 is shown. Number of high confident somatic variant identified in each adenoma is indicated on the right. A: adenoma; AC: adenocarcinoma; HGD: high grade dysplasia; CRC: colorectal cancer; wt: wild-type. Signatures with a contribution of less than 5% were merged into "other".

#### Supplemental Figure 1

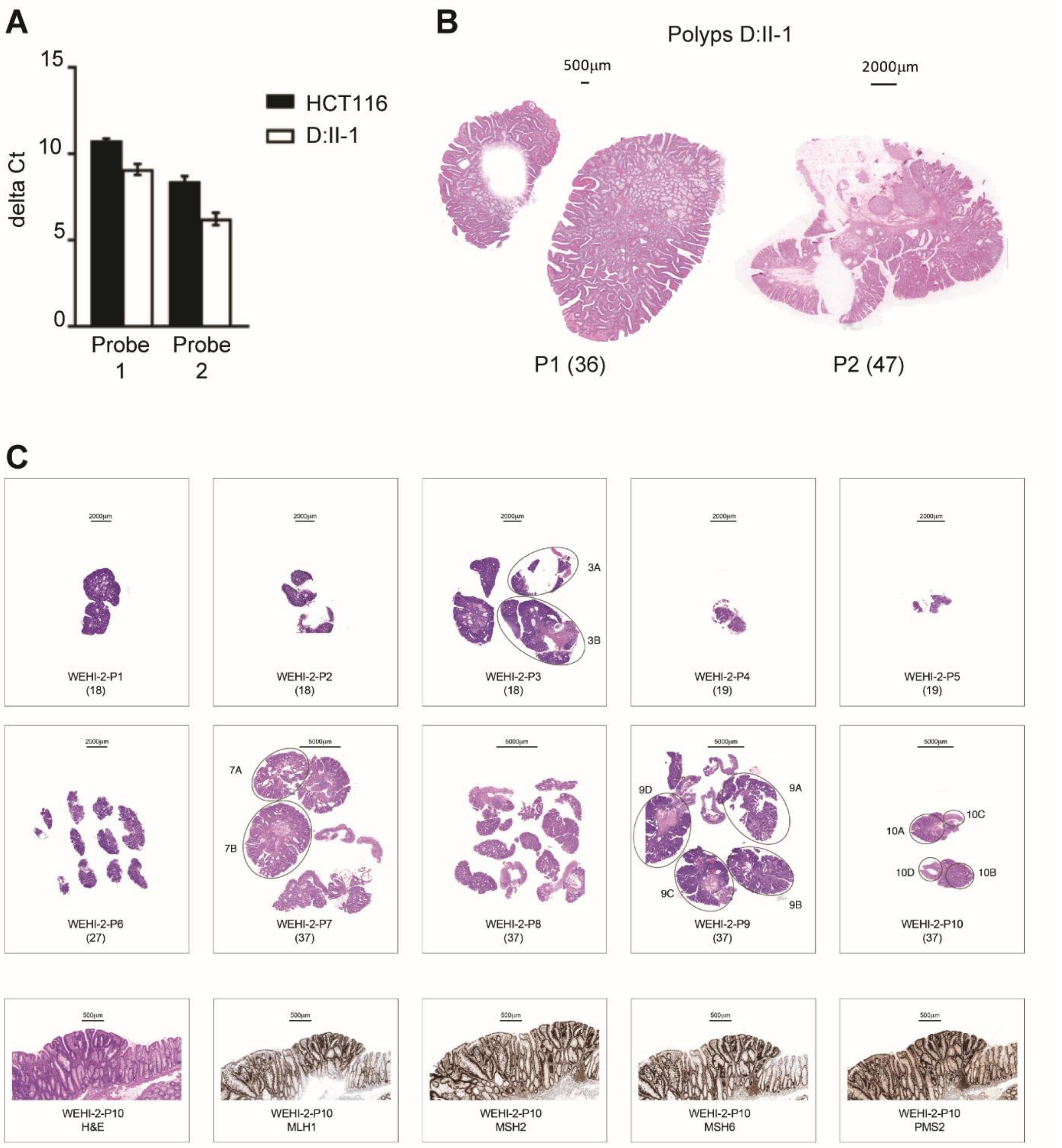

#### Supplemental Figure 2

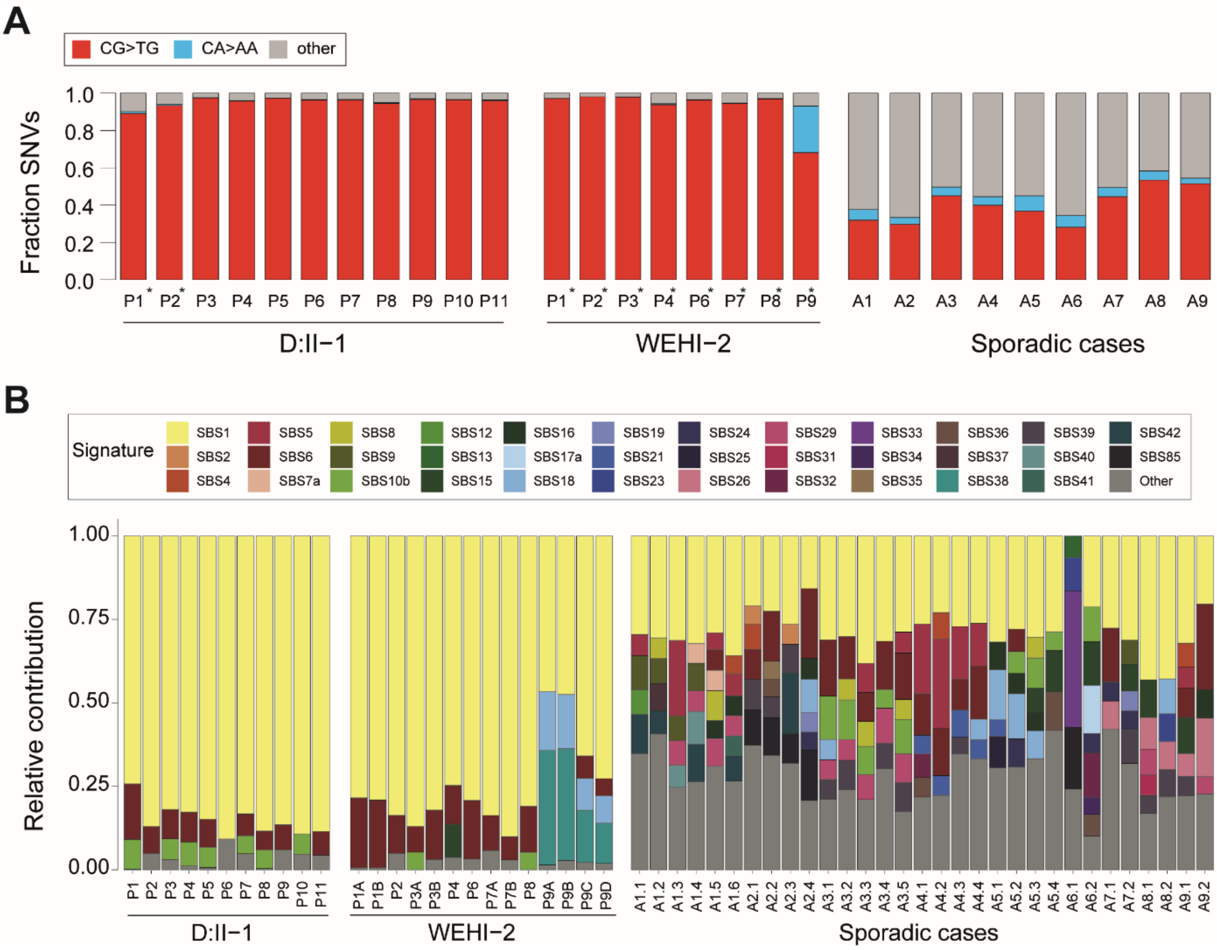

#### Supplemental Figure 3

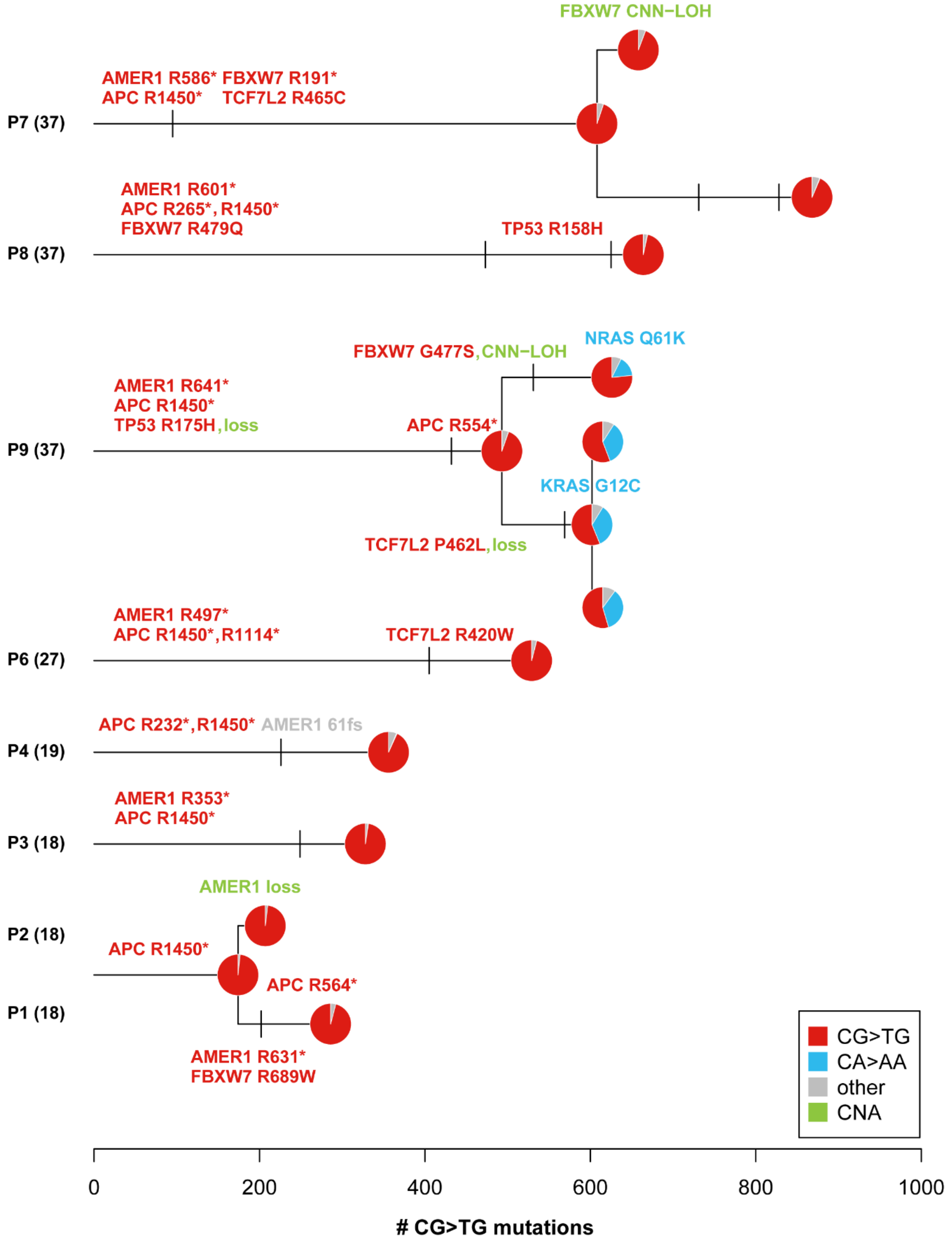

#### Supplemental Figure 4

**A**

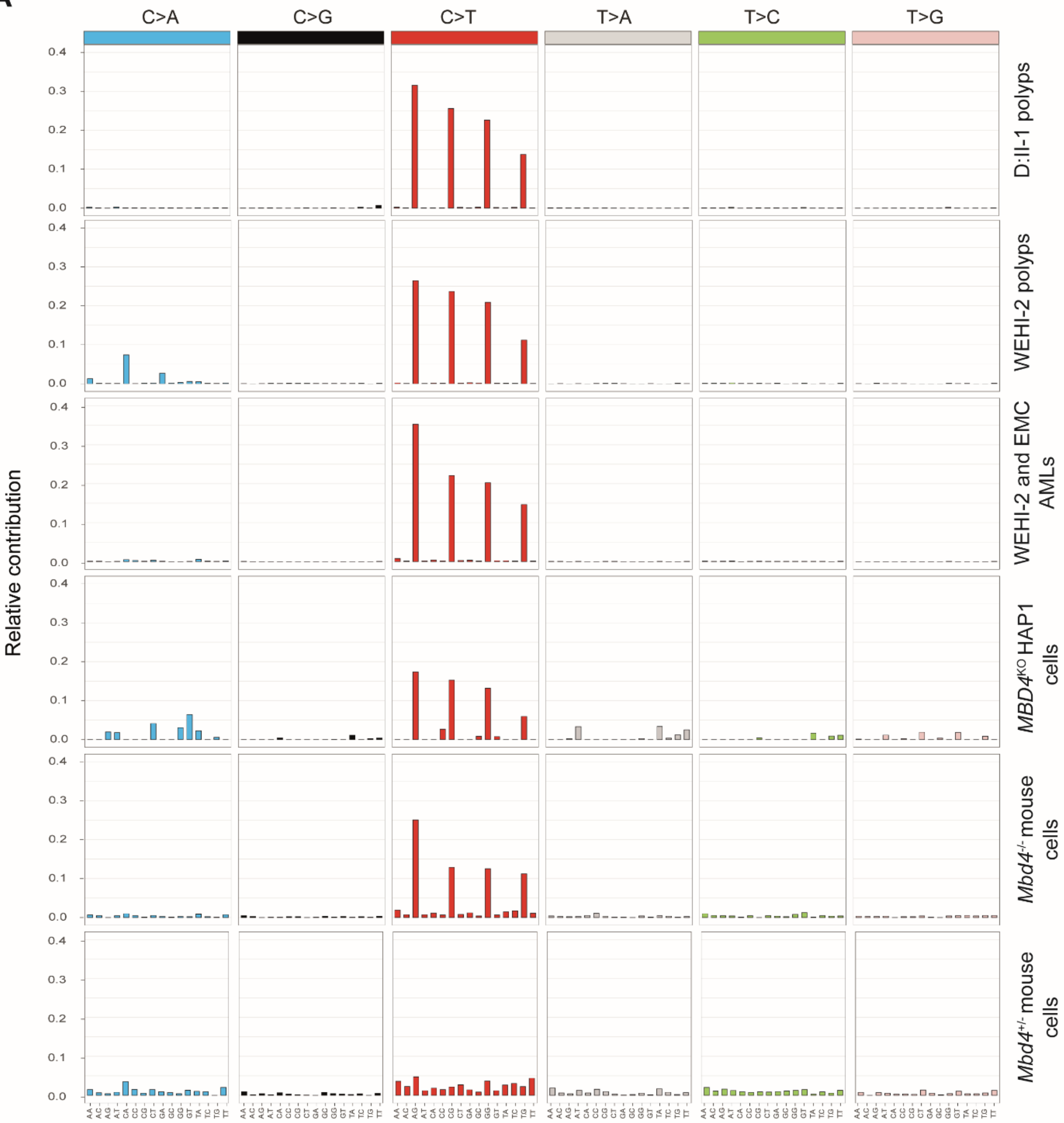


**B**

| Mutational profiles | D:II-1 | WEHI-2 |
|---|---|---|
| SBS1-v3 (COSMIC) | 0.9836 | 0.9602 |
| WEHI and EMC AMLs | 0.9928 | 0.9708 |
| MBD4KO HAP1 cells | 0.9683 | 0.9466 |
| MBD4 -/- mouse cells | 0.9193 | 0.9094 |
| MBD4 +/- mouse cells | 0.4867 | 0.5262 |

#### Supplemental Figure 5

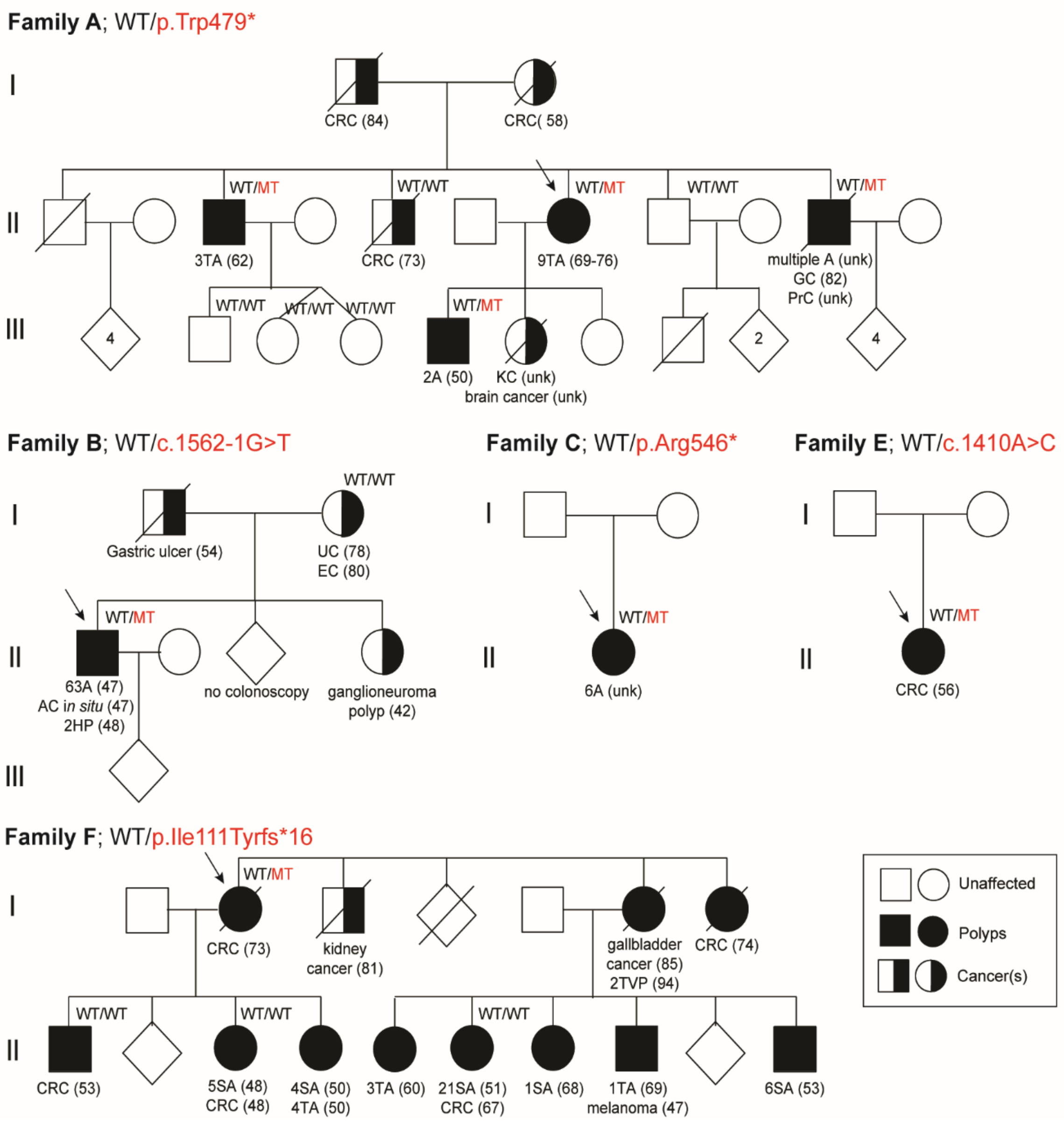

### Supplemental Figure 6

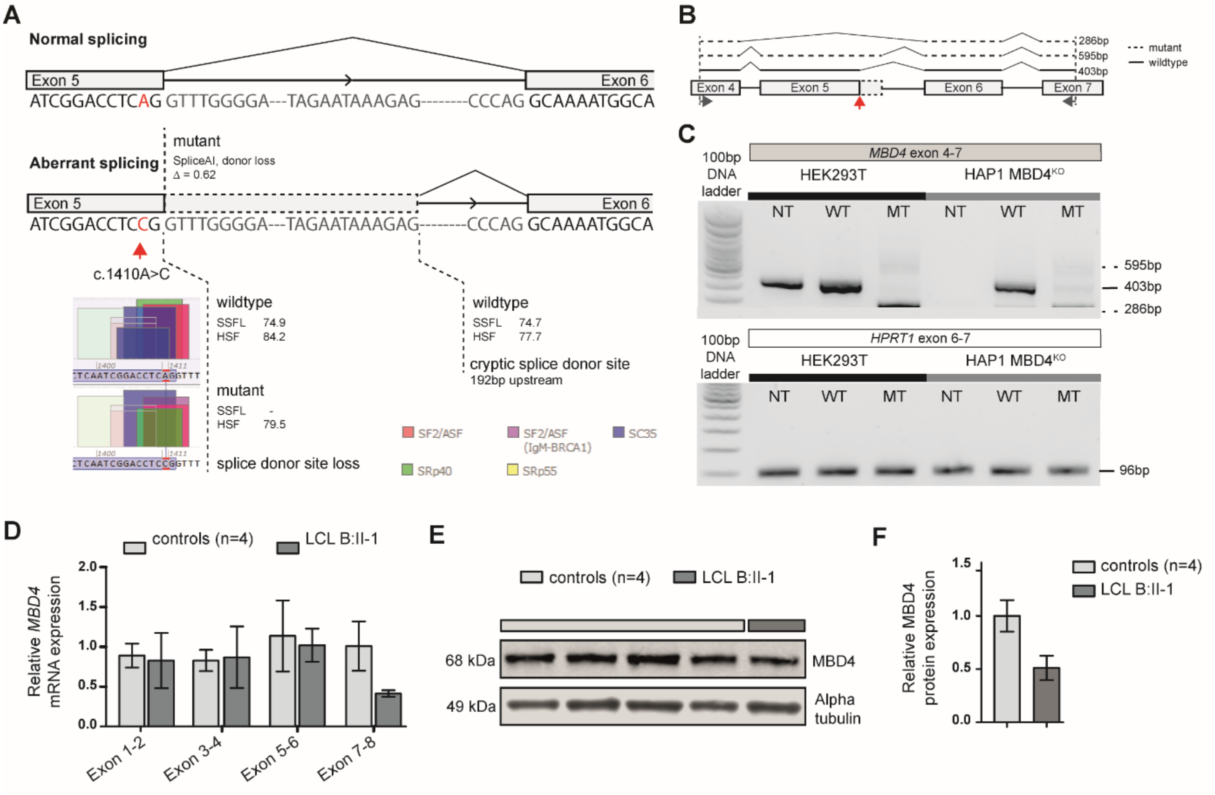


### Supplemental Figure 7

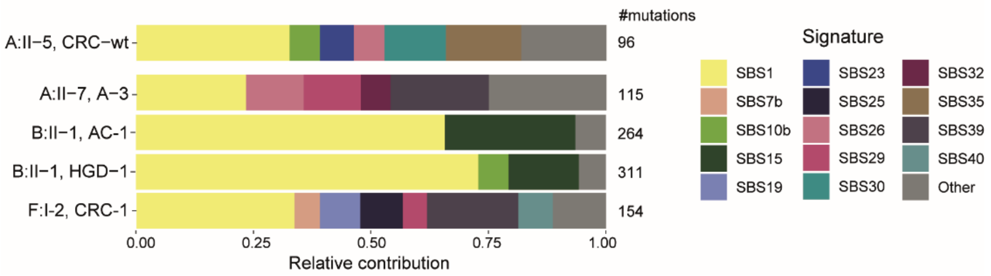
